# G51D mutation of the endogenous rat *Snca* gene disrupts synaptic localisation of α-synuclein priming for Lewy-like pathology

**DOI:** 10.1101/2023.10.27.564027

**Authors:** Stephen West, Ammar Natalwala, Karamjit Singh Dolt, Douglas J. Lamont, Melanie McMillan, Kelvin Luk, Tomoji Mashimo, Tilo Kunath

**Author notes:** Authors contributed equally (Stephen West, Ammar Natalwala). Corresponding authors: **Dr Stephen West:** **Professor Tilo Kunath:**. **Competing interests:** All authors have declared no competing interests.

## Abstract

Point mutations in the *SNCA* gene, encoding α-synuclein (αSyn), are a known cause of familial Parkinson’s disease. The G51D mutation causes early onset neurodegeneration with complex pathology. We used CRISPR/Cas9 in rats to introduce the G51D mutation into the endogenous *Snca* gene. Co-localisation immunostaining studies with synaptic proteins showed that αSyn^G51D^ protein is no longer efficiently localised to synapses. Furthermore, biochemical isolation of synaptosomes from rat cortex demonstrated a significant depletion of αSyn in *Snca^G51D/+^* and *Snca^G51D/G51D^* rats. Unbiased proteomic investigation of the cortex identified significant synaptic dysregulation in *Snca^G51D/G51D^* animals. Finally, we compared the propensity for Lewy-like pathology of *Snca^+/+^* and *Snca^G51D/G51D^* rats by stereotaxically delivering αSyn pre-formed fibrils (PFFs) into the pre-frontal cortex. At an early time-point, 6 weeks post-injection, we observed discrete Lewy-like structures positive for phosphoserine-129-αSyn (pS129-αSyn) only in *Snca^G51D/G51D^* brains. At 26 weeks post-injection of PFFs *Snca^G51D/G51D^* brains exhibited intense, discrete pS129-αSyn-positive structures, while *Snca^+/+^* brains exhibited diffuse pS129-αSyn immunostaining. Quantification of discrete pS129-αSyn-positive structures revealed the striatum of *Snca^G51D/G51D^* rats had significantly more Lewy-like pathology than *Snca^+/+^* rats. In summary, this novel *Snca^G51D^* rat model exhibits molecular characteristics of early synaptic dysfunction and is primed for αSyn pathology.

## Introduction

Most Parkinson’s disease cases are sporadic, and a relatively small number of cases are caused by *SNCA* multiplication or point mutations (Singleton *et al*., 2003; Chartier-Harlin *et al*., 2004). The latter include the A30G, A30P, E46K, G51D, A53E, A53G, A53T, A53V and E83Q mutations (Polymeropoulos *et al*., 1996; Krüger *et al*., 1998; Zarranz *et al*., 2004; Pasanen *et al*., 2014; Yoshino *et al*., 2017; Kapasi *et al*., 2020; Liu *et al*., 2021). The G51D point mutation, in exon 3 of *SNCA*, was discovered in 2013, and reported in a French, British and later a Japanese family (Kiely *et al*., 2013; Lesage *et al*., 2013; Tokutake *et al*., 2014). It is of particular interest, as the clinical manifestation of this mutation was that of early-onset and severe features of Parkinson’s disease, dementia with Lewy bodies (DLB), and multiple system atrophy (MSA), with clinical severity comparable to patients with a *SNCA* triplication mutation (Singleton *et al*., 2003). Furthermore, glial cell inclusions were observed in G51D patient brains on autopsy. The frontal cortex, hippocampus and posterior striatum were sites of observed pathology, in addition to the substantia nigra (Kiely *et al*., 2015). The accelerated disease progression in G51D patients makes this an important mutation to study and to delineate the underlying pathological mechanisms of conditions linked with α-synucleinopathy.

Limited animal models incorporating the G51D mutation have been generated so far, and whilst data suggests that this mutation leads to dopaminergic cell loss and diminished survival in *Drosophila* (Mohite *et al*., 2018), this was not the case in a pig model (E46K, H50Q and G51D mutations combined, with H50Q no longer being regarded as pathogenic) (Zhu *et al*., 2018). We generated a rat model of *Snca^G51D^* using a CRISPR/Cas9 approach (Morley *et al*., 2023). This overcomes limitations of other animal models incorporating transgenic overexpression, such as unknown transgene integration sites, variable transgene expression and pleiotropic effects (Nuber *et al*., 2013). We recently reported positron emission tomography data for our G51D rat model, showing an increase in dopamine turnover in the striatum in aged *Snca^G51D/G51D^* rats (Morley *et al*., 2023). However, there was no loss of dopamine production. This may suggest that in the rat, there are compensatory mechanisms mitigating the severity of dopamine loss observed in patients with the G51D mutation.

The first aim of this study was to characterise αSyn expression and its subcellular localisation in *Snca^G51D/G51D^* and *Snca^+/+^* rats. Based on the biophysical properties of αSyn^G51D^ and its reluctance to form α-helices (Fares *et al*., 2014), we hypothesised mis-localisation of αSyn from the synapse. Secondly, we aimed to investigate if there is any dopaminergic cell loss and the presence of Lewy-like pathology. Finally, we show that pathology can be readily triggered in *Snca^G51D/G51D^* and *Snca^+/+^* rats using human wild-type αSyn pre-formed fibrils (PFFs) injected stereotaxically into pre-frontal cortex, with mutant animals revealing an accelereated pattern of disease propagation.

## Materials & Methods

### Animals

All animal experiments were conducted under the Project Licence No. PC6C08D7D in accordance with Home Office regulations under the Animals Scientific Procedures Act (ASPA) 1986. Rats used in this study were maintained on a Fischer 344 background and were either wild-type at the *Snca* locus or were heterozygous or homozygous for the *Snca^G51D^* mutant allele (Morley *et al*., 2023). Rats used in this study were given *ad libitum* access to food and water and were maintained on a 12-hour light-dark cycle. All rats used were culled by schedule 1 CO_2_ inhalation followed by decapitation.

### Genotyping

Genotyping was performed as previously described (Morley *et al*., 2023). Briefly, genomic DNA was extracted from ear notches and *Snca* exon 3 was PCR amplified (forward primer 5’-TGGTGGCTGTTTGTCTTCTG-3’ and reverse primer 5’-TCCTCTGAAGACAATGGCTTTT-3’). The G**GA** to G**AT** (G51D) mutation created a new *Bsp*HI site in exon 3. PCR products were subject to *Bsp*HI restriction enzyme digest and agarose gel electrophoresis to distinguish wild-type, *Snca^G51D/+^* and *Snca^G51D/G51D^* mutant rats.

### Tissue dissection

Following schedule 1 cull, brains were removed, and depending on the downstream use, the brain was prepared using the following techniques:

1. Brain regions were sub-dissected and frozen on dry ice for synpatosome isolation, RNA isolation (for qRT-PCR), αSyn ELISA, or protein isolation for western blotting.
2. Brain was submerged in 4% PFA/PBS, for 24 hours followed by further processing for immunohistochemical analysis.

### RNA isolation

A pestle and mortar were cooled in a dry ice ethanol bath and used to crush tissue to powder. RNA was extracted from ∼10 mg of tissue per animal. RNA was extracted using RNeasy Lipid kit (Qiagen, 74804), following manufacturer’s instructions. RNA was quantified using the Qubit 2.0 (ThermoFisher) system following manufacturers instruction’s. RNA quality was assessed using the TapeStation2200.

### qRT-PCR

RNase-free water (Ambion, AM9937) was added to the RNA (200 ng - 1 μg) to give a 11 μl sample. The samples were incubated with 1 μl dNTP mix (10 mM, Life Technologies, 10297018) and 1 μl random primers (50 ng/μl, Thermo Fisher, PCR-545-020T) at 65**°**C for 5 minutes and then chilled on ice. After brief centrifugation, 4 μl 5x SSIV Buffer (ThermoFisher, 18090010), 1 μl 0.1M DTT (Life Technologies, Y00147) and 1 μl RNAse OUT (40 units/μl, Life Technologies, 10777019) were added. Then 1 μl Superscript IV reverse transcriptase (ThermoFisher, 18090010) or 1 μl RNase free water for the negative reverse transcriptase sample was added. This was mixed and incubated at room temperature for 10 minutes, followed by 10 minutes at 53**°**C. The reaction was inactivated by incubation at 80**°**C for 10 minutes. The cDNA mix was placed on ice and 60-180 μl RNase free water was added. The cDNA samples were then ready for qPCR.

qPCR was performed using the Roche LightCycler® 480 System with the Universal Probe Library (UPL) (Roche). The Roche UPL Assay design centre was used to design intron-spanning primers (IDT) with a specific UPL probe (Table 1). Reactions containing primers, UPL Probe, UPL Probe Master Mix (Roche) and PCR water were performed in 386-well plates as described in the manufacturer’s instructions. The results were normalised to the geometric mean of two house-keeping genes (*Tbp*, *Hprt*) and then expressed relative to control samples.

**Table 1:**
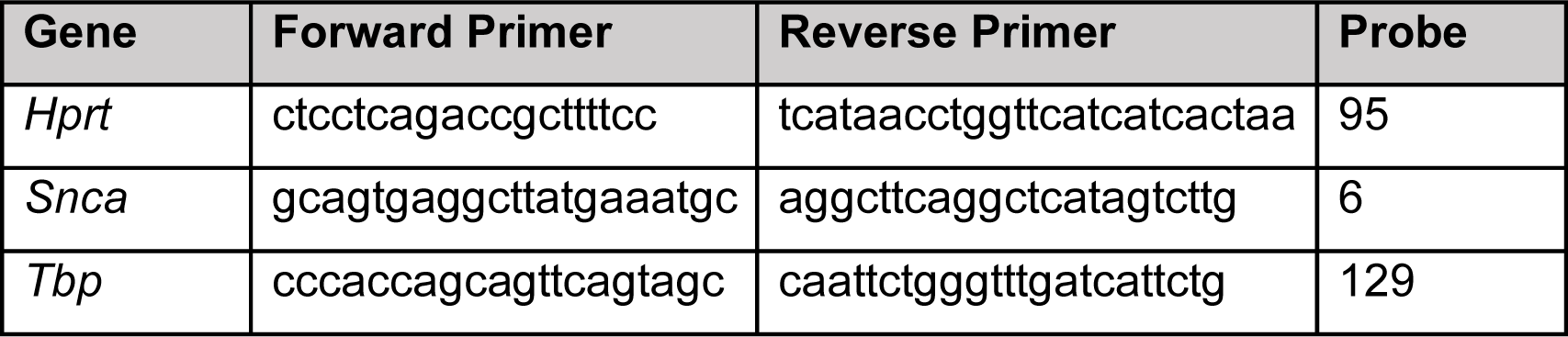
Primers and Probes used in UPL qPCR system αSyn ELISA.

ELISA for αSyn protein (Bioassay Technology Laboratory, E1113Ra) was performed as manufacturer’s instructions. Optical density (OD) was determined in FLUOstar Omega (BMG LABTECH) plate reader set to 450nm. Line of best fit is fitted to standards, and equation of the line is generated. This equation is used to calculate protein concentration of the samples from their OD.

### DAB Immunohistochemistry

Following 24-hour PFA fixation, tissue was washed 3xPBS (Gibco, 18912014) over 24 hours. At this point tissue was either bisected and prepared for sagittal sections (αSyn) or sub-dissected into regions of interest using a rat brain matrix (TH analysis). Regions were chosen based on the Rat Brain Atlas (Paxinos and Watson, 2013). The most anterior point of the cortex (bregma +5mm) was used as a zero point. For striatal analysis 3mm to 6.5mm was dissected (bregma +2 to −1.5mm) and for nigral analysis 9mm to 12.5mm (bregma −4.5mm to −7.5mm) was dissected. Dissected regions were then cryoprotected in 30% sucrose (SLS, CHE3650)/PBS (Gibco, 18912014) until the tissue sunk (usually 24-48 hrs) at 4°C. Tissue was then frozen on a metal chuck with optimum cutting temperature (OCT, CellPath, KMA010000A) compound in a dry ice ethanol bath and stored at −20°C until sectioning. Every 6^th^ 30um serial section was taken using the Leica cryostat (Cyrostar NX50) set at −20°C onto Superfrost slides (ThermoFisher, J1800AMNT).

Following sectioning slides were washed in TBS, and then blocked in 10% goat serum (Sigma, G9023)/TBS with 0.1% Triton (Fisher, BP151-100) (10% GS/TBT) for 1 hour. Slides were then incubated overnight in rabbit anti-TH (1/1000) or rabbit anti-αSyn (1/1000) in 10% GS/TBT at 4°C in a humidity chamber. Slides were then washed 2 x 15 mins in TBT, and 1 x 15mins in TBS. Slides were then incubated with 0.3% hydrogen peroxide (Sigma H1009)/TBS to quench endogenous peroxidases for 15 mins. Slides were then rinsed in tap water, before incubation with a goat anti-rabbit IgG HRP-conjugated antibody (1/2000) for 30 mins. Slides were then washed 2 x 15 mins in TBT, and 1 x 15 mins in TBS, followed by incubation with the ImmPACT^TM^ DAB substrate (Vector Labs, SK-4105), for 5-10 mins. Tissue sections were imaged on the Axioscan Z.1 (Zeiss) slide scanner, using a Plan-Apochromat 20x/NA 0.8 Objective. Light source intensity was set at 221% and pixel size was 0.22 μm x 0.22 μm. Adaptive focus was set using the Onion Skin 53 strategy. Images were captured on the Hitachi HV-F202SCL with a 200 μs exposure time.

Striatal tyrosine hydroxylase images were analysed in QuPath v0.4 (Bankhead *et al*., 2017). Briefly, a region of interest was placed around the striatum and cortex, the optical density of DAB staining in the cortex was considered background and subtracted from the staining in the striatum. The optical density across the A-P axis of the striatum was averaged for each animal. The staining was normalised to control animals and compared across genotypes. Nigral tyrosine hydroxylase images were analysed using the cell counter to count TH+ cell bodies in the substantia nigra. The cell counts across the A-P axis of the nigra was averaged for each animal. Counts were normalised to control animals and compared across genotypes.

DAB immunohistochemistry for pSer129-αSyn was done at 6 weeks with the same manual protocol. At 6 months, the sections were processed on the Leica Bond Max. These brains were embedded in paraffin and sectioned. The paraffin sections were de-waxed and re-hydrated using xylene and graded alcohols. Antigen retrieval was done using Epitope retrieval buffer 1 (pH 6; Leica biosystems) for 20 minutes. pS129-αSyn primary antibody (AB51253, 1:500) was used. Bond wash with TBS buffer (Fischer scientific) and Tween 20 was used for washing. Briefly, the protocol was: wash 10 minutes, peroxide block 5 minutes, wash, primary antibody applied for 60 minutes, wash, polymer 15 minutes, bond wash and wash with distilled water, DAB immunostain applied for 10 minutes, wash with distilled water, Haematoxylin stain 5 minutes and final wash. Reagents were from Leica Bond Polymer Refine detection kit (DS 9800). DAB images of αSyn PFF-injected brains were scanned with Axioscanner (Zeiss) using a 20x objective NA 0.8 Plan Apochromat. QuPath open source software was used to quantify pSer126-αSyn pathology using a custom script (Bankhead *et al*., 2017).

### Fluorescent Immunohistochemistry

Tissue was prepared as above, but 10 μm sections were taken instead of 30 μm. Slides were brought up to room temperature, and then microwaved for 20 secs in TBS to remove OCT. Slides were then blocked in 10% goat serum in TBS with 0.1% Triton (10% GS/TBT) for 1 hour. Slides were then incubated overnight in rabbit anti-αSyn (1/1000) and mouse anti-synaptophysin (1/50) in 10% GS/TBT at 4°C in a humidity chamber. Slides were then washed 3 x 15 mins in TBT. Slides were then incubated with goat anti-rabbit IgG Alexa Fluor 488 and goat anti-mouse IgG2a Alexa Fluor 568 (both 1/1000) in 10% GS/TBT for 2 hours at room temperature. Slides were then washed 3 x 15 mins in TBT, followed by 5-minute incubation with DAPI/TBS (1:10000), and 3 x 5 mins TBS washes. Slides were then mounted using ProLong^TM^ Diamond Antifade Mountant (Thermo Fisher, P36965) and 1.5 high precision coverslips (Marienfeld, 0107172).

### Confocal microscopy and image analysis

Images were acquired on a Leica TCS SP8 5D confocal microscope. Images were taken at 63x magnification using a HC PL APO 63x/1.4 oil objective, with a refraction index of 1.518 2x Zoom. Pinhole was set to 1AU. Images were acquired at a speed of 600 hz. Line Accumulation was set at 14. Line Sequential scanning was performed and for DAPI, the 405 nm laser was set at 2.9%. HyD detector was used to capture the emitted photons in counting mode, and gain was set at 75. For αSyn immunofluorescence detection the 488nm laser was set at 2.99%. HyD detector was used to capture the emitted photons in counting mode, and gain was set at 70. For synaptophysin immunofluorescence detection the 594nm laser was set at 20%. HyD detector was used to capture the emitted photons in counting mode, and gain was set at 89. Images were deconvolved with the Huygens (SVI) software using an CMLE algorithm, with 40 iterations and a signal-to-noise ratio set to 10. Deconvolved images were subsequently used for co-localisation analysis. The built-in Huygens (SVI) (pixel-based) co-localisation analyser was used to calculate Spearman’s correlation coefficient (r) values, and background estimation set to ‘optimised’.

### Cortical protein preparation and synaptosome Isolation

Brains were harvested from twelve 6-month-old rats (4 rats (3 male and 1 female) for each genotype *Snca^+/+^*, *Snca^G51D/+^* and *Snca^G51D/G51D^*). The frontal cortex was dissected from the rest of the brain and then bisected along the midline. Total protein was isolated from one half of the cortex and synaptosomes were isolated from the other half as previously described (Llavero Hurtado *et al*., 2017). Briefly, each half cortex was frozen in low-bind 2-ml Eppendorf tubes (Eppendorf, 0030108132) prior to addition of 500 μl of 0.32 M Sucrose Solution (0.32 M sucrose (SLS, CHE3650), 1 mM EDTA (Invitrogen, 15575-038), 5 mM Tris (Roche, 107089176009), pH 7.4). Each sample was homogenised until smooth and centrifuged at 900g for 10 mins at 4°C. Supernatants were transferred to new Eppendorf tubes and each pellet was resuspended in 500 μl 0.32 M Sucrose Solution. Each sample was spun at 900g for 10 mins at 4°C, and this supernatant was combined with supernatant from first spin. The pellet from the second spin was and considered the non-synaptic fraction. The combined supernatant samples were spun at 20000g for 20 mins at 4°C, supernatant was discarded and the pellets were the synaptosome fractions. See Supplementary Figure 1 for overview. All samples were stored at −80°C until protein extraction.

### Western blotting

Protein concentrations were determined using the Micro BCA Protein Assay Kit (Thermo Scientific, 232350), following the manufacturer’s instructions. Protein (15 μg) was incubated with 5 μl NuPAGE® Loading Buffer (4x, Life Technologies, NP0008), 2 μl 1 M DTT and made up to 20 μl with deionised water at 100°C for 5 mins. The gel (4-12% Bis-Tris NuPAGE® gel, Life Technologies, NP0322BOX) was loaded into the tank with running buffer (20x NuPAGE® MOPS SDS Running Buffer in deionised water, Life Technologies, NP0001). Protein samples were loaded along with the SeeBlue Plus 2 protein marker (LC5925, Life Technologies). The gel was run at 60V for 20 mins, and then at 120V until the blue loading dye reached the base of the gel (usually 90 mins) at room temperature. Gels were removed from case and then processed for Coomassie staining (see below) or transferred for immunostaining.

PDVF membranes (Amersham Hybond ECL, GE Healthcare Life Science, RPN68D) were activated in methanol for 30 seconds and then incubated with transfer buffer (3.02 g Tris (Roche, 107089176009), 14.4 g glycine (Sigma, G7126), 800 ml water and 200 ml methanol (Fisher, M/3900/17)). The gel was removed and prepared for transfer to PDVF membrane. It was then loaded into the transfer tank, transfer buffer was added and the blot was run at 300 mA for 75 mins at 4°C.

The membrane was removed dried for 30 mins, then reactivated in methanol (30 seconds). Membrane was then incubated in blocking solution (5% milk in 0.01% TBT) for 2 hrs at RT. After removal of blocking solution, the membrane was incubated with primary antibody in blocking solution over-night at 4°C. The membrane was washed 3 times for 15 mins in 0.01%TBT and then incubated with appropriated secondary antibody (anti-mouse HRP; 1:2000, or anti-rabbit HRP; 1:2000) in blocking buffer for 1 hour at room temperature. Subsequently, the membrane was washed in 0.01%TBT 3 times for 15 mins before HRP was detected using the Pierce ECL Western Blot substrate kit (Thermo Scientific, 32109), according to manufacturer’s instructions. Membrane was placed in an acetate cassette and then imaged on the LI-COR Odyssey® Fc, using the 700nm channel (30 sec exposure) for ladder, and Chemi channel (10-minute exposure) for antibody of interest. LI-COR Image Studio^TM^ was used to analyse images, box (size maintained between samples) was placed around band and OD was measured within box. This was then normalised to loading control OD (analysed in same way). Samples were expressed as % of control samples.

### Coomassie staining and pooling samples

Following gel run, gel was removed from case and incubated in Coomassie stain (50%v/v Methanol (Fisher, M/3900/17), 10%v/v acetic acid (SLS, CHE1014), 0.2% Coomassie Brilliant blue R-250 (Sigma, B7920)) for 30 mins. Gels were then de-stained by 4 x 1hour washes in De-stain solution (7.5%v/v methanol, 10%v/v glacial acetic acid). Gels were then placed in an acetate cassette and imaged on the LI-COR Odyssey® Fc, using the 700nm channel and exposed for 10 mins. Using LI-COR Image StudioTM, OD of staining was determined for each sample at 4 weights, full lane 28-62 kDa, 62-198 kDa and 2-28 kDa. Following this, samples for TMT-Mass spectrometry were pooled, and 50 μg of each sample was added to respective pool and made up to 100 μl with label free buffer. Coomassie gels were run again and analysed by the same method to check equivalency of loading.

### Quantitative mass spectrometry

Samples for proteomics was performed by the Fingerprints Facility, University of Dundee. Sample preparation and protein identification and quantification analysis by mass spectrometry was carried out as previously described (Llavero Hurtado *et al*., 2017). MaxQuant Version 1.6.0.16 was used for the assignment of peptides to protein using the uniport-rat-jan2018.fasta file. A min and max peptide length of 8 and 25 aa respectively was used for unspecified search. The mass spectrometry proteomics data have been deposited to the ProteomeXchange Consortium via the PRIDE partner respository (Perez-Riverol *et al*., 2022) with the data set identifier PDX0044776.

### Pairwise comparisons and pathway analysis

Following filtering to remove non-unique and unquantified peptides, pairwise comparisons were made between the different mutant samples and *Snca^+/+^* for both the cortical and synaptosome samples. In all cases Log2(Fold Change) compared to *Snca^+/+^* was ranked in order (Supplementary Table 1), and proteins were taken forward for pathway analysis if their expression was changed 20% up (Log2(FC)>0.26) or down (Log2(FC)<-0.32) relative to *Snca^+/+^*. Protein lists generated by pairwise analysis were uploaded to g:profiler (http://biit.cs.ut.ee/gprofiler/gost), an online tool which performs functional enrichment analysis, mapping genes to known functional information sources and detects statistically significantly enriched pathways (Raudvere *et al*., 2019). Organism was set to *Rattus norvegicus* and significance threshold was set at 0.05 based on the g:SCS algorithm. We identified enriched biological pathways using the Kyoto Encyclopaedia of Genes and Genomes (KEGG) (Kanehisa *et al*., 2019) (Supplementary Table 2).

### Stereotactic injection of human αSyn^WT^ pre-formed fibrils

Rats were anaesthetised using isofluorance per standard protocol. Depth of anaesthesia checked using tail pinch and blink reflex. Each rat was positioned on the stereotaxy frame (Stoelting, 51950), hair shaved, scalp skin cleaned with iodine and 2% Lidocaine administered subcutaneously. A clear drape was applied. Lacrilube to both eyes. Under sterile conditions, a vertical midline scalp incision (1 cm) was performed and skin retracted using retractors (Stoelting, 52124). Bregma was located using the digital arm manipulator and marked with a shallow burr hole (Stoelting, 51449V). Target co-ordinates (AP 3.2 ML −1.5 DV −2.5) were used to place burr hole. A 10 µl Hamilton syringe (Stoelting, 51105), pre-filled with human wild-type αSyn PFFs (10 µg in 4 µl) (Luk *et al*., 2012), was used to penetrate the dura and to reach the target co-ordinates for injection. The αSyn PFFs were injected at 1 µl/minute rate with the Quintessential Stereotaxic Injector and the needle left in situ for two minutes. This was done to allow αSyn PFFs to settle locally and the needle slowly retracted to minimise reflux. The skin was sutured with dissolvable 4-0 vicryl. Post-operatively, rats were placed on a heat pad at 30°C (Stoelting, 53850-RR) until motor recovery in a clean cage. The wound was checked and rats weighed regularly. Mash food and analgesia in drinking water (Rimadyl 5 mg/kg) and Vetergesic jelly 0.5 mg/kg was given in the post-operative period.

### Statistical analysis

Where data fell into a normal distribution, Graph Pad Prism v.6.0c was used to generate a parametric one-way ANOVAs with Tukey’s multiple comparisons test to determine statistical significance of differences between *Snca^+/+^*, *Snca^G51D/+^* and *Snca^G51D/G51D^* animals. In the case of analysis just comparing *Snca^+/+^* and *Snca^G51D/+^* or *Snca^+/+^* and *Snca^G51D/G51D^* then unpaired two-tailed Student’s t-test was used. If data was not normally distributed, a non-parametric Kruskal-Wallis test with Dunn’s multiple comparisons test was performed to determine statistical significance of differences between *Snca^+/+^*, *Snca^G51D/+^* and *Snca^G51D/G51D^* animals. In the case of analysis just comparing *Snca^+/+^* and *Snca^G51D/+^* or *Snca^+/+^* and *Snca^G51D/G51D^* animals, then a non-parametric unpaired two-tailed Mann Whitney test was used. See individual figure legends for details. P<0.05 was considered significant.

## Results

### G51D mutagenesis of *Snca* does not significantly alter the overall abundance of αSyn protein in the rat brain

The introduction of a GGA-to-GAT (G51D) codon change into the endogenous rat *Snca* gene did not affect embryonic or fetal development, and progeny from heterozygous matings were born in Mendelian ratios (Morley *et al*., 2023). We first explored whether the G51D mutation impacted on the level of αSyn expression within the brain. qRT-PCR was performed using RNA extracted from the brainstem of *Snca^+/+^* and *Snca^G51D/G51D^* rats and no significant change in *Snca* mRNA levels was observed at 12 months (Figure 1A). Further, total αSyn protein levels, measured using ELISA, from the cortex of the *Snca^+/+^* and *Snca^G51D/G51D^* rats at 12 months was not significantly different (Figure 1B). DAB immunohistochemistry for total αSyn showed similar total and regional expression in *Snca^+/+^* and *Snca^G51D/G51D^* rat brains across multiple regions in both male and female rats (Figure 1C,D). The comparable appearance of αSyn immunostaining between genotypes, in each of the brain regions investigated, supports the *Snca* qRT-PCR and αSyn ELISA data.

**Figure 1.**
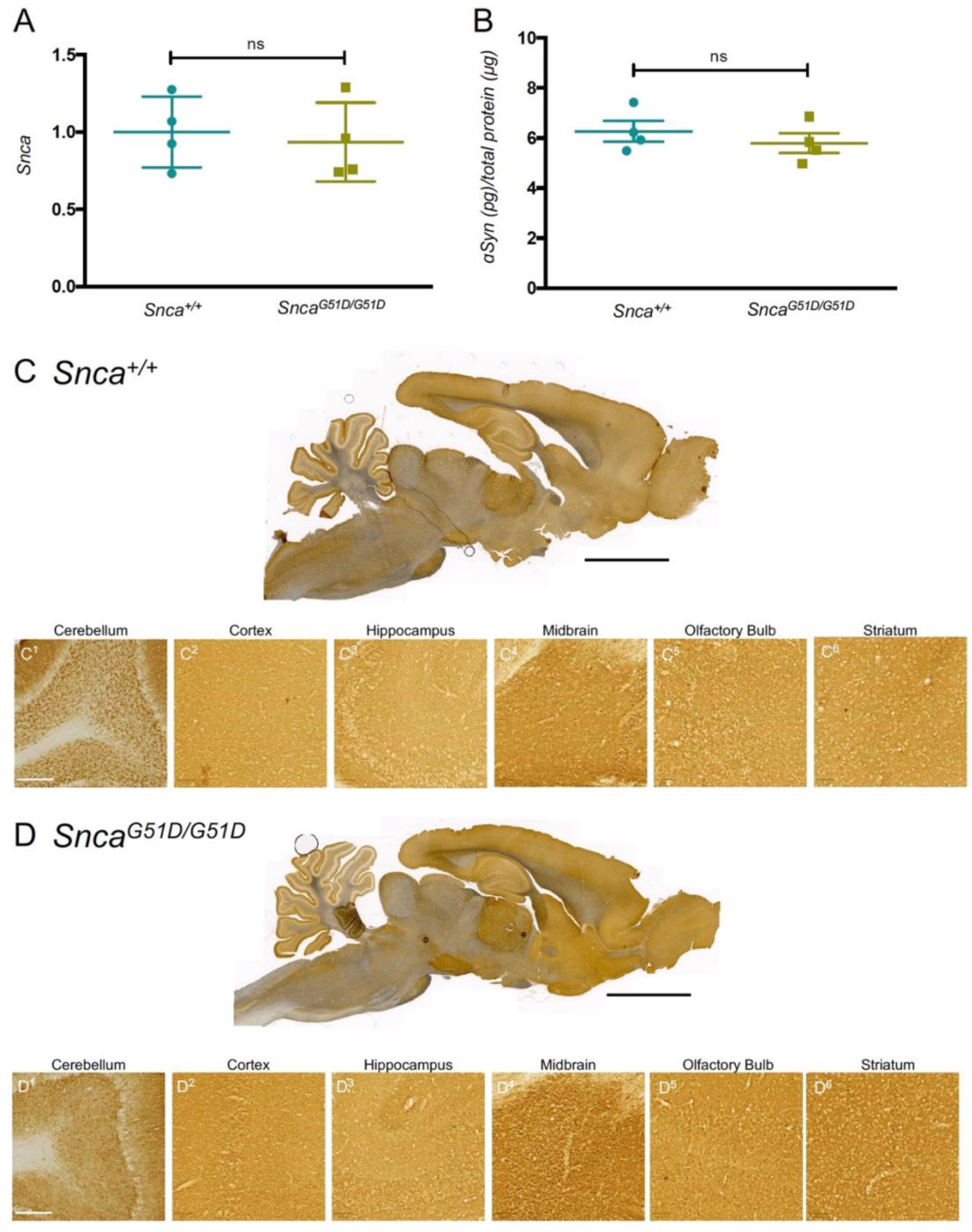
*Snca* transcript levels and αSyn protein amounts are not altered by the G51D mutation. αSyn expression across multiple brain regions appear normal in the *Snca^G51D/G51D^* rat brain. **A.** qPCR showing *Snca* gene transcript levels from the brain stem relative to the arithmetic mean of *Hprt* and *Tbp* and normalized to controls. There is no significant difference in transcription levels of *Snca* in *Snca^+/+^* and *Snca^G51D/G51D^* 12-month-old animals, (n=4 (2 males, 2 females) rats per genotype; p=0.72, unpaired t-test). Data presented as mean±SD. **B.** ELISA measuring αSyn concentration in the cortex relative to total protein. There is no difference (p=0.44, unpaired t-test) in αSyn abundance in the cortex of *Snca^G51D/G51D^* (5.78±0.79 pg/μg) compared to controls (6.27±0.83 pg/μg) at 12 months of age, n=4 (2 males, 2 females) animals per genotype. Data presented as mean±SD. **C,D.** DAB immunohistochemistry for αSyn in sagittal sections. Scale bar 5 mm. There are no differences in αSyn levels between genotypes and no inclusion formation in the *Snca^G51D/G51D^* brains. C^1-6^, D^1-6^: High magnification of different brain regions. Scale bar, 200 μm.

### αSyn^G51D^ is mis-localised from the synapse in *Snca^G51D/G51D^* rat brain

Given that abundance of αSyn protein was not significantly changed in *Snca^G51D/G51D^* rats, we next examined whether there was any change in the subcellular localisation of αSyn. Thus, sections of cortex from *Snca^+/+^* and *Snca^G51D/G51D^* rats were immunostained for αSyn and the pre-synaptic marker, synaptophysin. Image analysis of αSyn and synaptophysin localisation in the cortex indicated that there was a significant decrease in the amount of co-localisation between these two proteins in the *Snca^G51D/G51D^* animals compared to *Snca^+/+^* controls (Figure 2A-C).

**Figure 2.**
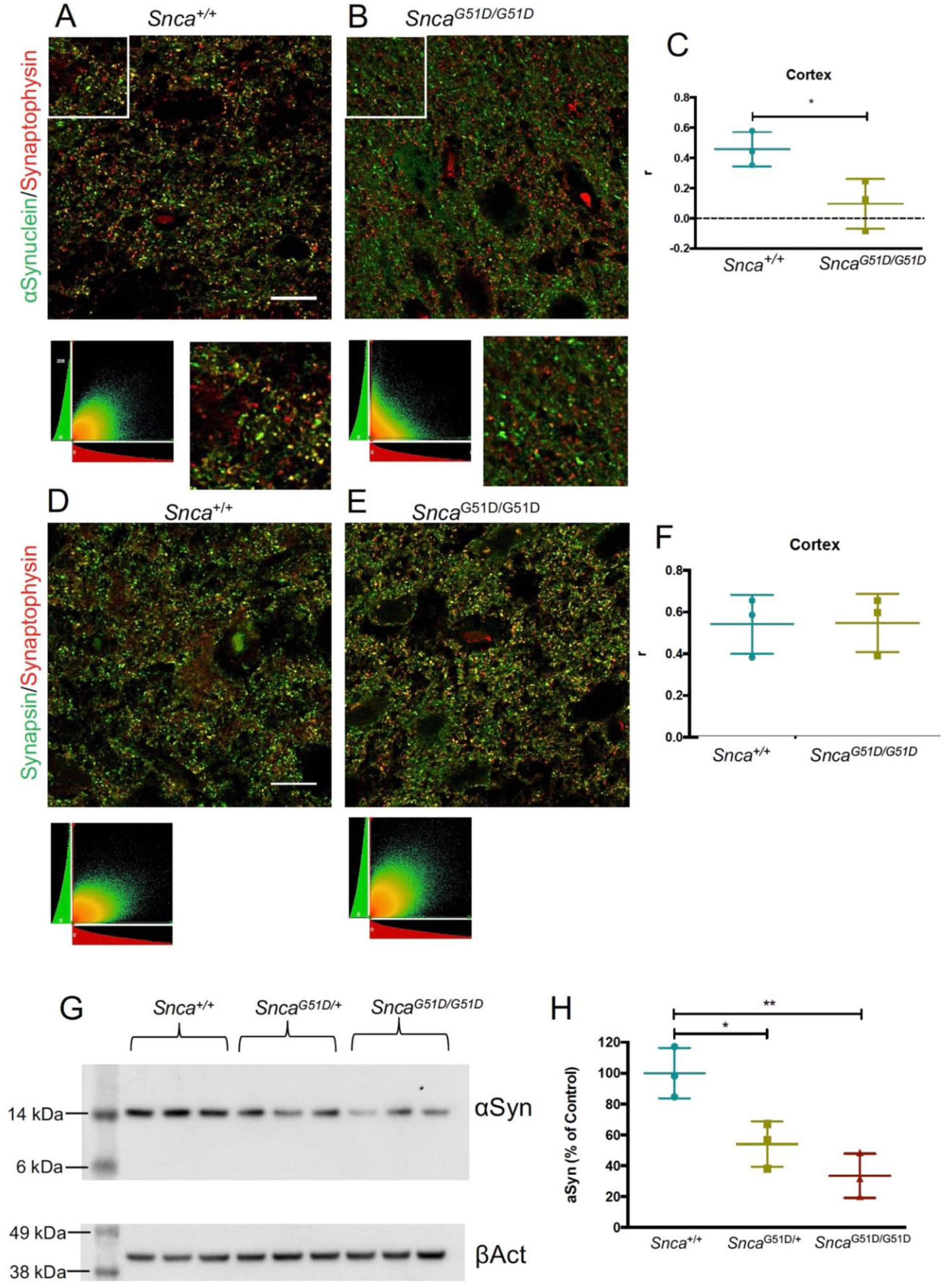
αSyn is depleted from synapses in *Snca^G51D/G51D^* rat cortex. **A,B.** Representative deconvolved confocal images showing co-localisation between αSyn and synaptophysin, a pre-synaptic marker. Below confocal image is co-localisation plot showing pixel intensity in green channel (y-axis), plotted against pixel intensity in red channel (x-axis) and inset images are zoom of white boxes. **C.** Comparison of object Spearman’s (r) values for different genotypes. There was a significant decrease in the average r value between *Snca^+/+^* and *Snca^G51D/G51D^* in the cortex (C, *Snca^+/+^*=0.458±0.11, *Snca^G51D/G51D^*=0.095±0.17, p=0.035, unpaired two-tailed t-test). **D,E.** Representative deconvolved confocal image showing co-localisation between synapsin and synaptophysin, two pre-synaptic markers. Below confocal image is co-localisation plot showing pixel intensity in green channel (y-axis), plotted against pixel intensity in red channel (x-axis) D = *Snca^+/+^* and E = *Snca^G51D/G51D^*. **F.** Comparison of object Spearman’s (r) values for different genotypes. There was no difference between the average r value between *Snca^+/+^* (0.54±0.14) and *Snca^G51D/G51D^* (0.58±0.14) suggesting that synaptic structure is maintained in the *Snca^G51D/G51D^* animals. Data presented as mean±SD, n=3 (2 males, 1 female) animals per genotype. Scale bar 15 μm. **G**. Western blot for αSyn in synaptosome samples of all genotypes, with b-Actin loading control. **H**. Quantification of αSyn levels in **G**, relative to β-actin and normalized to *Snca^+/+^*. There is a significant effect of genotype (p=0.0045, one-way ANOVA) on the levels of αSyn in the cortical synaptosomes of *Snca^+/+^*, *Snca^G51D/+^* (54.0%±14.5) and *Snca^G51D/G51D^* (33.6%±14.3) rats at 6 months and multiple comparison analysis showed significant differences between *Snca^+/+^*and *Snca^G51D/+^* (p<0.05) and *Snca^+/+^* and *Snca^G51D/G51D^* (p<0.01), n=3 (2 males and 1 female).

To confirm that this was due to a change in the localisation of αSyn and not due to a disruption in synaptic structure, another pre-synaptic marker, synapsin, was co-labelled with synaptophysin. In both *Snca^+/+^* and *Snca^G51D/G51D^* rats there was a positive and similar level of co-localisation of these synaptic markers (Figure 2D-F). Taken together, this suggests that *Snca*^G51D/G51D^ rats have no major disruption of overall synaptic structure, but contain significantly reduced amounts of αSyn protein.

To further investigate the synaptic depletion of αSyn by a complementary method, we performed western blot analysis for αSyn on isolated synaptosome samples (Figure 2G). There was a significant difference in αSyn levels between genotypes at 6 months. Multiple comparison analysis showed a significant reduction in αSyn between *Snca^+/+^* and *Snca^G51D/+^* (p<0.05) and *Snca^+/+^*and *Snca^G51D/G51D^* (p<0.01) groups (Figure 2H).

### Proteomic analysis showed dopaminergic synapse and Parkinson’s disease pathway dysregulation in *Snca^G51D/G51D^* rat cortex

Brains were harvested from four (4) 6-month old rats of each genotype (3 male and 1 female) and the frontal cortex was dissected from the rest of the brain and then bisected along the midline. Total protein was isolated from one half and the other half was spun through sequential sucrose gradients to isolate the synaptosomes before protein was extracted (Supplementary Figure S1A) (see Materials and Methods for further detail). Western blotting for the H2A histone family member X (H2AX) demonstrated the absence of the nuclear protein from the synaptic fraction (Supplementary Figure S1B). The whole cortex and synaptosome samples of all three genotypes (*Snca^+/+^*, *Snca^G51D/+^*, *Snca^G51D/G51D^*) were processed for tanden mass tag (TMT) mass spectrometry at the Fingerprints Proteomics Facility at the University of Dundee. A total of 8211 unique proteins were identified across all samples. The data set was filtered for proteins identified with at least 2 unique peptides and were observed in all six conditions, which left a total of 6640 proteins for downstream analysis. Proteins that were more than 20% up-regulated (66 proteins) or more than 20% down-regulated (79 proteins) in *Snca^G51D/G51D^* relative to *Snca^+/+^* cortex (Figure 3A, Supplementary Table 1), were used for KEGG pathway analysis. There were 13 significantly enriched KEGG pathways for the dysregulated proteins including Parkinson’s disease (KEGG:05012), dopaminergic synapse (KEGG:04728) and oxidative phosphorylation (KEGG:00190) (Supplementary Table 2). The disease pathways Parkinson’s disease and Alzheimer’s disease in the *SNCA^G51D/G51D^* cortex were significant due to down-regulation of proteins involved in oxidative phosphorylation (NDUFAB1, COX5A, NDUFB8), and in the case of PD dopamine transport (SLC18A2 or VMAT2).

**Figure 3.**
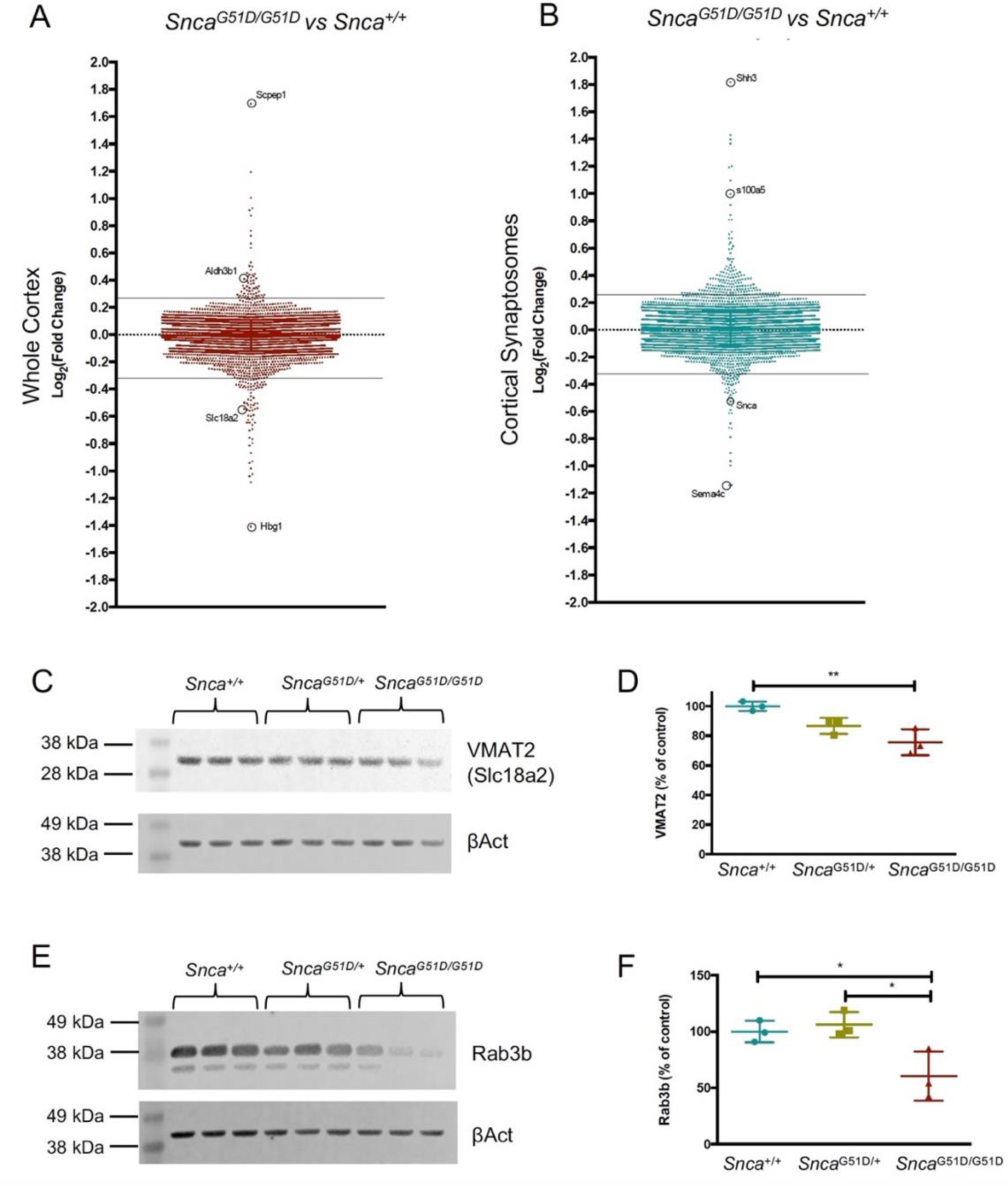
Proteomic analysis of whole cortex and cortical synaptosomes. **A.** Log2(Fold Change) of all proteins in *Snca^G51D/G51D^* compared to *Snca^+/+^* cortex. Solid lines represent cut off for 20% up-regulated (>0.26) or down-regulated (<-0.32). **B.** Log2(Fold Change) of all proteins in *Snca^G51D/G51D^* compared to *Snca^+/+^* cortical synaptosome. See Supplementary Table S1 for lists of proteins. **C.** Western blot for VMAT2 in cortical samples, with β-actin loading control. **D.** Quantification of VMAT2 levels in **C**, relative to β-actin and normalized to *Snca^+/+^*. There is a significant effect of genotype (p=0.0087, one-way ANOVA) on VMAT2 expression in the cortex of *Snca^+/+^*, *Snca^G51D/+^* (86.7±5.4%) and *Snca^G51D/G51D^* (75.7±8.8%) rats at 6 months and multiple comparison analysis show a significant difference between *Snca^+/+^*and *Snca^G51D/G51D^* (p<0.01). n=3 (2 males and 1 female) per genotype. **E**. Western blot for Rab3b in synaptosome samples, with β-actin loading control. **F**. Quantification of Rab3b levels in **E,** relative to β-actin and normalized to control. There is a significant effect of genotype (p=0.02, one-way ANOVA) on the levels of Rab3b in the cortical synaptosomes of *Snca^+/+^*, *Snca^G51D/+^* (106.1±11.2%) and *Snca^G51D/G51D^* (60.6%±21.9%) rats at 6 months and multiple comparison analysis show significant differences between *Snca^+/+^*and *Snca^G51D/G51D^* (p<0.05) and *Snca^G51D/+^* and *Snca^G51D/G51D^* (p<0.05). n=3 (2 males and 1 female) per genotype for both analyses.

### *Snca^G51D/G51D^* synaptosomes analysis reveals dysregulation of serotonergic synapse and actin cytoskeleton

Proteins that were either 20% up-regulated (163 proteins) or down-regulated (60 proteins) in the *Snca^G51D/G51D^* synaptosome compared to *Snca^+/+^* (Figure 3B, Supplementary Table 1) were used for KEGG pathway analysis. There were 5 significantly enriched KEGG pathways when the dysregulated proteins were analysed (Supplementary Table 2). Spliceosome (KEGG:03040), serotonergic synapse (KEGG:04726), and regulation of actin cytoskeleton (KEGG:04810). Serotonergic innervation is known to be affected in early PD (Kerenyi *et al*., 2003), therefore it is expected to see this pathway being enriched in the *Snca^G51D/G51D^* synapses.

### VMAT2 is down-regulated in the *Snca^G51D/G51D^* cortex

Validation of data from the TMT-mass spectrometry experiment was performed using western blot analysis for key dysregulated proteins. Of these, VMAT2 was selected as it is known to be canonically involved in Parkinson’s disease (Pifl *et al*., 2014). VMAT2 was down-regulated in *Snca^G51D^*(log_2_ (Fold Change) −0.54 in *Snca^G51D/+^*and −0.55 in *Snca^G51D/G51D^*) relative to *Snca^+/+^*cortex. By western blotting we observed a dose-dependent decrease in the levels of VMAT2 protein in the cortex across the genotypes (Figure 3C,D), and there was a significant downregulation between *Snca^G51D/G51D^* and *Snca^+/+^* cortex (p<0.01; Figure 3D).

### Rab3b is down-regulated in *Snca^G51D/G51D^* cortical synaptosomes

We chose to validate the synaptosome protein Rab3b (log_2_ (Fold Change) −0.39 in *Snca^G51D/+^* and - 0.30 in *Snca^G51D/G51D^*) compared to *Snca^+/+^,* which when over-expressed in the rat substantia nigra has been shown to protect dopaminergic neurons in the 6-OHDA lesion model (Chung *et al*., 2009). There was a significant reduction in Rab3b in the synaptosomes of each gentype at 6 months, and in *Snca^G51D/G51D^* compared to *Snca^+/+^* rats (p<0.05; Figure 3E,F), in agreement with the mass spectrometry data.

### Tyrosine hydroxylase-positive neurons within the substantia nigra do not degenerate in aged *_SncaG51D/G51D_* _rats_

Following the finding of the loss of αSyn from the synapse, we aimed to characterise the viability of the dopaminergic system in each genotype. In Parkinson’s, αSyn aggregation occurs throughout the brain, yet dopaminergic neurons are particularly susceptible to degeneration (Petrucci *et al*., 2016). In late stages of PD, degeneration is also observed in cortical areas which can lead to dementia; this is especially the case in familial PD caused by the αSyn G51D mutation (Kiely *et al*., 2013; Lesage *et al*., 2013). Quantification of TH^+^ neurons indicated there was no difference between the number of TH^+^ neurons in the substantia nigra of *Snca^+/+^*, *Snca^G51D/+^* and *Snca^G51D/G51D^* animals at either 6 months or 12 months (Supplementary Figure S3A-H). This suggests that in this rat model, αSyn^G51D^ does not cause overt death of dopaminergic neurons by 12 months of age. Quantification of the optical density of TH staining in the striatum, showed that there was no difference between genotypes at 6 months (Supplementary Figure S3I-L). This highlights that there was no down-regulation of TH expression within the striatum of the mutant animals.

### *Snca^G51D/G51D^* brains are more susceptible to Lewy-like pathology following human wild-type αSyn PFF injection

Human wild-type αSyn pre-formed fibrils (PFFs) were used to trigger disease pathology in *Snca^+/+^* and *Snca^G51D/G51D^* animals via intracerebral injection into the pre-frontal cortex. DAB immunostaining for pSer129-αSyn was used to detect Lewy-like pathology and while no inclusions were seen in *Snca^+/+^* controls, at six weeks post PFF injection there were Lewy-like structures in the pre-frontal cortex of *Snca^G51D/G51D^* rats near the injection site and the olfactory bulb (Figure 4A). At six months, following pre-frontal αSyn PFF injection, both *Snca^+/+^* and *Snca^G51D/G51D^* rats developed Lewy-like inclusions, however, the overall abundance was greater at six months and different in appearance between the genotypes. G51D homozygous mutants had well-defined inclusions that appeared to be denser and more intensely stained, whereas control rats had diffuse, nuclear and cytoplasmic pSer129-αSyn staining. The striatum, specifically, had more Lewy-like pathology in *Snca^G51D/G51D^* rats than controls (Figure 4B-C). These data suggest that rats harbouring αSyn^G51D^ are highly susceptible to acquiring Lewy-like pathology and areas such as the striatum are specifically vulnerable to pathology in this model of α-synucleinopathy.

**Figure 4.**
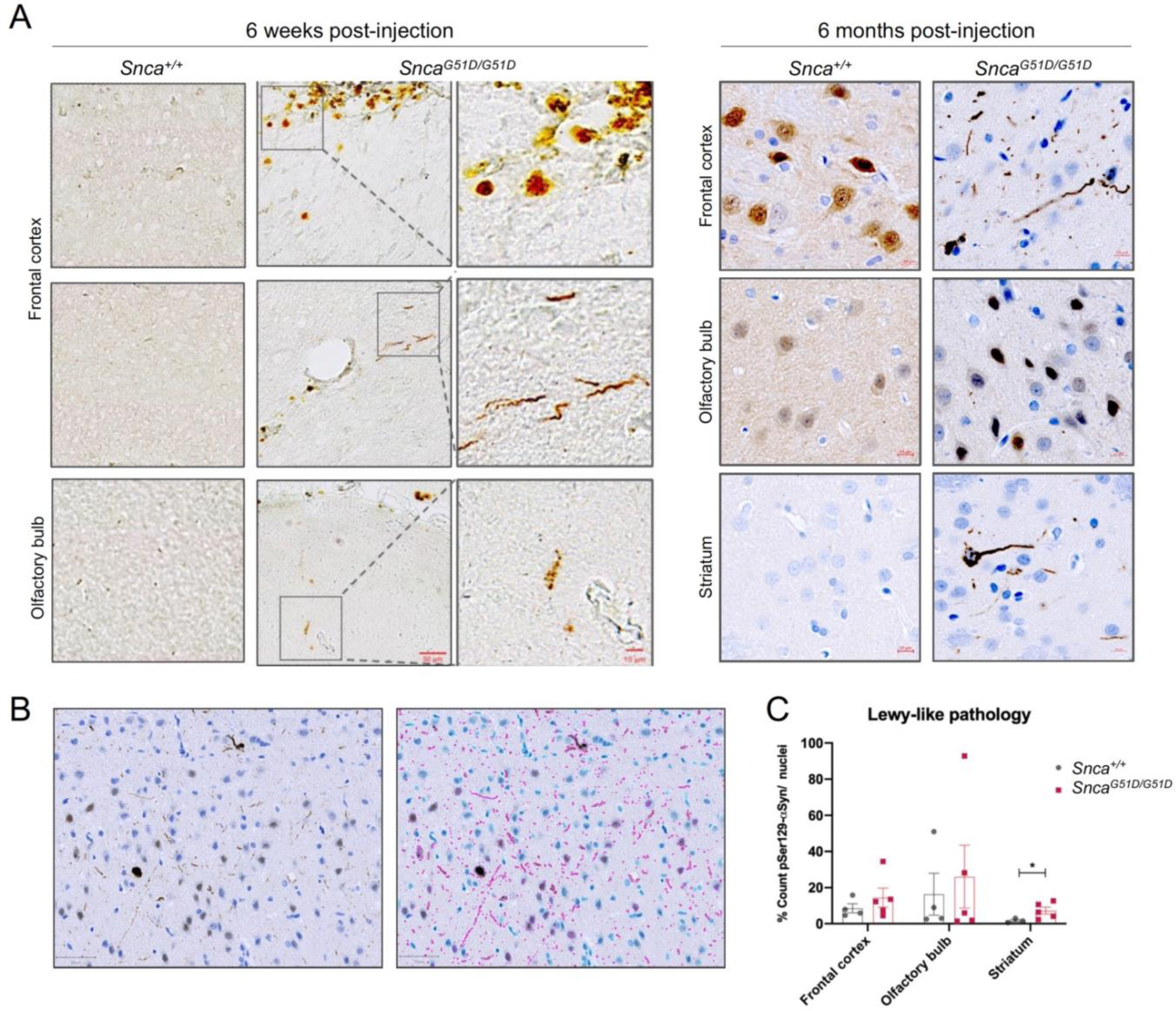
*Snca^G51D/G51D^* rats are primed for pathology and develop Lewy-like structures preferentially in the striatum at 6 months following human αSyn^WT^ PFF injection. **A.** pSer129-αSyn DAB immunostaining in adult *Snca^+/+^* and *Snca^G51D/G51D^* rats at 6 weeks and 6 months post-injection of human αSyn^WT^ PFFs into the prefrontal cortex. At 6 weeks, Lewy-like structures formed near the injection site in the *Snca^G51D/G51D^* rat brain, but absent from *Snca^+/+^* rat brain. Scale bar, 50 µm (10 µm, zoomed-in image). The 6-month images include haematoxylin (blue) nuclear staining and show Lewy-like pathology in the olfactory bulb, frontal cortex and striatum. Scale bar, 10 µm. These inclusions were diffuse in *Snca^+/+^* compared to well-defined and mature appearing ones in *Snca^G51D/G51D^* rats. **B.** Images to show detection of nuclei and Lewy-like structures, using a custom script in QuPath. **C.** Quantification of Lewy-like pathology per nuclei at 6 months showed significantly more inclusions in the striatum of *Snca^G51D/G51D^* rats than controls (*Snca^+/+^*, 1.73 ± 1.19; *Snca^G51D/G51D^*, 7.21 ± 4.34; p<0.05, t-test).

## Discussion

In this study, we characterised a novel rat model of α-synucleinopathy generated by CRISPR/Cas9 engineering to create an endogenous *Snca^G51D^* mutation (Morley *et al*., 2023). The G51D mutation alone was insufficient to cause the formation of Lewy-like pathology and nigrostriatal degeneration at one year. However, we identified that αSyn was mis-localised away from the synapse in homozygous *Snca^G51D/G51D^* rats, priming for pathology once a disease trigger, specifically intracerebral αSyn PFFs, is introduced. Unbiased proteomic analysis revealed subcellular dysfunction at the level of the synapse and mitochondria, as well as aberration within Parkinson’s disease-related pathways. This work provides an excellent model to investigate αSyn-mediated disease *in vivo*, with several advantages over αSyn transgenic overexpression models, and could provide major insights into the early disease mechanism occurring in α-synucleinopathies.

In comparison to the *SNCA* triplication mutation, where αSyn levels are increased two-fold (Devine *et al*., 2011), we show that both RNA and protein quantification for total αSyn was similar in homozygous *Snca^G51D/G51D^* mutant and control rats. This suggests that the pathogenicity of the αSyn^G51D^ protein is more aggressive than αSyn^WT^ given that the clinical phenotypic severity of both mutations in patients is similar (Petrucci *et al*., 2016). We expected to observe Lewy-like pathology in the *Snca^G51D/G51D^* rats and subsequent altered levels of total αSyn, but this was not the case at one year. It is possible that with even older mutant rats we may have observed this pathology, but nonetheless, compared to patients with αSyn^G51D^ there was a lack of an early onset disease phenotype. This is further corroborated by data from the study by Zhu and colleagues, where *SNCA* mutations at E46K, H50Q and G51D in combination, in minipigs did not lead to Lewy-like pathology at three months, albeit the animals harbored heterozygous mutations and the study interval was relatively short (Zhu *et al*., 2018). In support of the lengthy times needed to observe rodent phenotypes, the BAC transgenic rat model of αSyn overexpression did not exhibit significant formation of proteinase K insoluble αSyn deposits until 16 months of age (Nuber *et al*., 2013).

*In vitro* biophysical studies have shown that αSyn^G51D^ can impair phospholipid membrane affinity (Ysselstein *et al*., 2015), and αSyn^G51D^ can form different strains of PFFs under different experimental conditions, as well as cross-seed with αSyn^WT (^Peduzzo *et al*., 2020; Sun *et al*., 2021^)^. Different pathogenic αSyn strains can cause variable phenotypes, shown using brain homogenates from PD and MSA patients that were used to trigger pathology in cells or in rodent brains, with MSA-derived fibrils being more potent disease triggers (Van der Perren *et al*., 2020). We found αSyn^G51D^ to be mis-localised from synapses, but this was insufficient for Lewy-like structure formation. Fares and colleagues observed few inclusions resulting from αSyn^G51D^ but found an increase in mitochondrial fragmentation in cultured neurons (Fares *et al*., 2014). Interestingly, our proteomic analysis of *Snca^G51D/G51D^* cortex showed synaptic and mitochondrial dysfunction. It is possible that synaptic and mitochondrial dysfunction precede inclusion formation, which was apparent in our model following a second disease trigger. The lack of Lewy-like inclusions may explain the lack of observed dopaminergic neuronal loss in *Snca^G51D/G51D^*rats at one year. Although it is important to note that αSyn is more abundant in cortical excitatory synapses than dopaminergic synapses (Emmanouilidou *et al*., 2016), cortical dysfunction is more challenging to assess in a rodent model than in patients experiencing cognitive decline due to αSyn^G51D^. Our data also showed spliceosome pathway dysfunction due to αSyn^G51D^ and one study has demonstrated that the components of the spliceosome are present and functional in dendrites and may provide an additional level of local translational control (Glanzer *et al*., 2005).

A secondary disease trigger was a crucial element of this work to unmask the disease process in *Snca^G51D/G51D^* rats. Although the frontal cortex is not a commonly selected location for intracerebral αSyn injection (Reyes *et al*., 2014; Osterberg *et al*., 2015; Hao *et al*., 2018; Dhillon *et al*., 2019; Bassil *et al*., 2020), it is relevant to the G51D pathology observed on post-mortem and relevant to other α-synucleinopathies such as MSA. Further, it was useful to explore subsequent temporospatial Lewy-like pathology in interconnected regions such as the striatum. In both genotypes, the pathology was avid at the injection site and structures in proximity including the olfactory bulbs, but there was a predilection for inclusions in the striatum of *Snca^G51D/G51D^* rats. The Lewy-like inclusions appeared well-defined and mature, relative to diffuse nuclear and cytoplasmic pSer129-αSyn immunostaining in wild-type rats. This suggests that there is an active disease process in the *Snca^G51D/G51D^* rats such that propagation of pathology occurs at a faster rate, with a predilection for PD-vulnerable structures such as the striatum, in *Snca^G51D/G51D^* rats.

Whilst αSyn deletion or overexpression does not impair neurogenesis and development of immature neurons (Vargas *et al*., 2014; Natalwala *et al*., 2022), αSyn may have a role at the synapse in aged mature neurons. One of the limitations of this work is that rats older than one year were not examined and that gender differences were not examined. Another limitation is that we do not investigate the oligomeric state of αSyn in either of the genotypes. If the synaptic and mitochondrial dysfunction are early events in the *Snca^G51D^* model, analysis of oligomeric αSyn using super-resolution microscopy may reveal the extent of the underlying pathological process in *Snca^G51D^* rats (Murata *et al*., 2021; Xu *et al*., 2022). Furthermore, from post-mortem data we know that glia including oligodendrocytes exhibit significant αSyn^G51D^ pathology (Kiely *et al*., 2013), and this remains an area to be explored in this rodent model. Caveats regarding αSyn^WT^ PFFs as a disease trigger include the fact that the exact concentration cannot be precisely measured and delivered into the brain and different strains have different pathogenicity. Indeed Hayakawa and team shows that αSyn^G51D^ PFFs were more potent inducers of Lewy-like pathology than αSyn^WT^ PFFs (Hayakawa *et al*., 2020), and thus may have shown a greater difference between then genotypes following intracerebral injection in *Snca^G51D^*rats.

In summary, this study characterized a novel rat model of synucleinopathy, harbouring an endogenous *Snca^G51D^* mutation, with physiological levels of αSyn and without the limitations encompassing animal models with ectopic transgenes. The combination of synaptic and mitochondrial dysfunction and the αSyn^G51D^ mis-localisation from the synapse may be early events in the disease process, increasing the propensity for overt disease following a second disease trigger. Our disease-relevant model provided an opportunity to establish the underlying mechanism of the *SNCA^G51D^* point mutation that exerts a severe clinical phenotype in patients. This may lead to new mechanistic insights and disease-modifying treatments for patients with conditions resulting from α-synucleinopathy.

## Acknowledgements

We would like to thank Dr Emma Lane for assistance in setting up intracranial stereotactic injections, Lyndsey Boswell for immunostaining, Prof Patrik Brundin for experimental advice, Dr Samatha Eaton & Prof Tom Wishart for advice and support of synaptosome isolation, William Mungall for rat husbandry and tissue collection, and the team in the Histology service at the Shared University Research Facilities (SuRF) at the University of Edinburgh.

