## Supplementary data for "G51D mutation of the endogenous rat *Snca* gene disrupts synaptic localisation of α-synuclein priming for Lewy-like pathology"

### Supplementary Figure S1

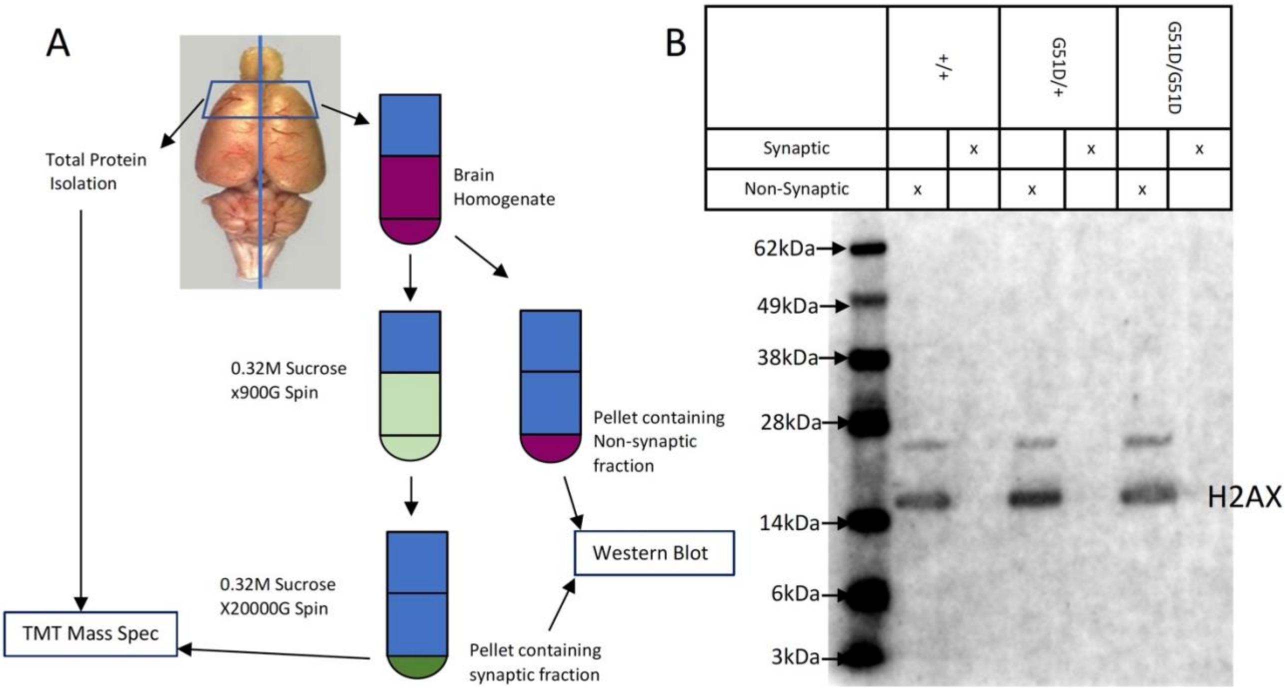


**Supplementary Figure S1. Whole cortex and synaptosome isolation for proteomic analysis. A.** Diagram outlining synaptosome and whole brain protein isolation for mass spectrometry. **B.** Western blot showing loss of nuclear protein H2AX from the synaptosome samples of all three genotypes.

**Supplementary Figure S2**

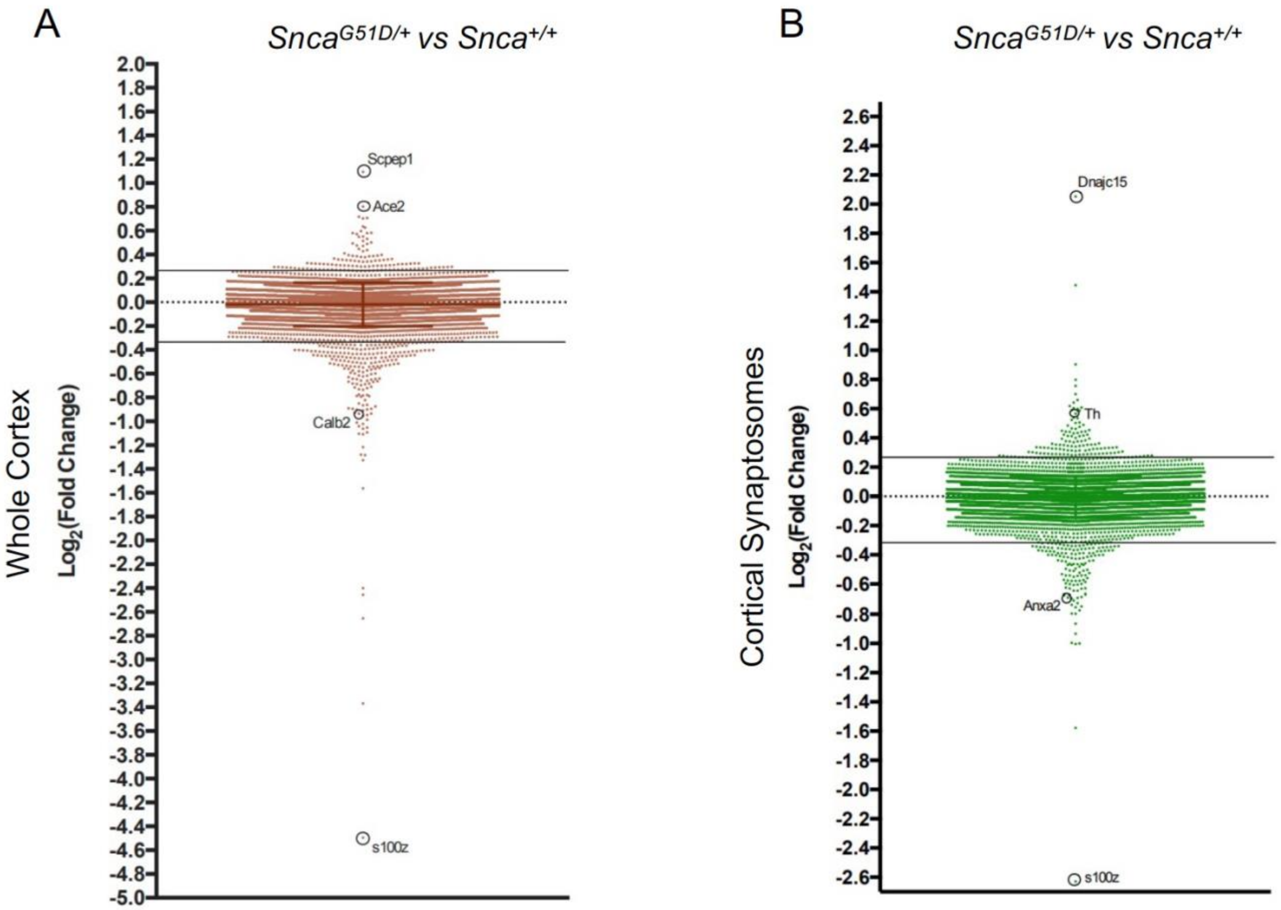


**Supplementary Figure S2. Proteomic analysis of *Snca*$^{G51D/+}$ whole cortex and cortical synaptosomes. A.** Log2(Fold Change) of all proteins in *Snca*$^{G51D/+}$ compared to *Snca*$^{+/+}$ cortex. Solid lines represent cut off for 20% up-regulated (>0.26) or down-regulated (<-0.32). (121 up-regulated and 251 down-regulated proteins). **B.** Log2(Fold Change) of all proteins in *Snca*$^{G51D/+}$ compared to *Snca*$^{+/+}$ cortical synaptosome preparations (134 up-regulated and 139 down-regulated proteins). See Supplementary Table S1 for lists of proteins.

**Supplementary Figure S3**

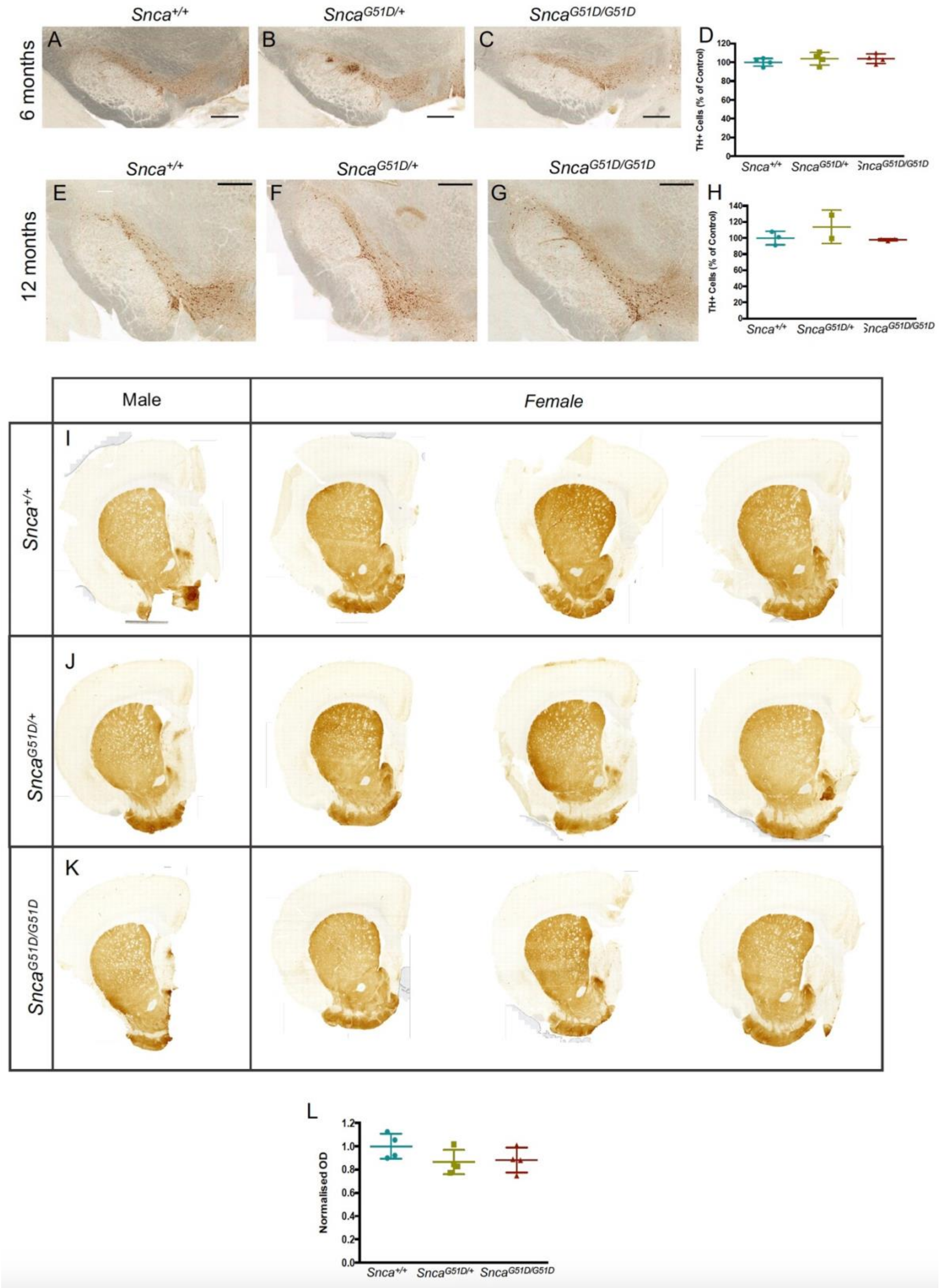


**Supplementary Figure S3. *Substantia nigra* TH+ neurons do not significantly degenerate in the *SNCA*$^{G51D/+}$ or *SNCA*$^{G51D/G51D}$ rats. A-C.** Representative images of TH+ neurons in the *substantia nigra* (SN) at 6 months. Scale bar, 500 µm. **D.** Quantification of

SN TH+ neurons in different genotypes relative to controls. There was no significant difference in cell numbers between *Snca*$^{+/+}$, *Snca*$^{G51D/+}$ and *Snca*$^{G51D/G51D}$ rats at 6 months. (n=4 (3 males, 1 female) animals per genotype; p=0.54, one-way ANOVA). **E-F.** Representative images of TH+ neurons in the SN at 12 months. Scale bar 500 µm D: Quantification of SN TH+ neurons in different genotypes relative to controls. There was no significant difference in cell numbers between *Snca*$^{+/+}$, *Snca*$^{G51D/+}$ and *Snca*$^{G51D/G51D}$ rats at 12 months. (n=3 males for *Snca*$^{+/+}$ and *Snca*$^{G51D/G51D}$, n=2 males for *Snca*$^{G51D/+}$; p=0.30, one-way ANOVA). I-K: Representative images of TH+ terminals in the striatum. I= *Snca*$^{+/+}$ J= *Snca*$^{G51D/+}$ K= *Snca*$^{G51D/G51D}$. Each coronal section is a representative slice for each animal analysed. Scale bar 1 mm. L: Quantification of striatal TH optical density (OD) in different genotypes relative to controls. There was no significant difference in OD between *Snca*$^{+/+}$, *Snca*$^{G51D/+}$ and *Snca*$^{G51D/G51D}$ rats at 6 months. (n=4 (3 males, 1 female) animals per genotype; p=0.20, one-way ANOVA). All data presented as mean±SD.
